# A fast, accurate, and generalisable heuristic-based negation detection algorithm for clinical text

**DOI:** 10.1101/2020.07.03.187054

**Authors:** Luke T Slater, William Bradlow, Dino FA Motti, Robert Hoehndorf, Simon Ball, Georgios V Gkoutos

**Affiliations:** College of Medical and Dental Sciences, Institute of Cancer and Genomic Sciences, University of Birmingham; Institute of Translational Medicine, University Hospitals Birmingham, NHS Foundation Trust; NIHR Experimental Cancer Medicine Centre; NIHR Surgical Reconstruction and Microbiology Research Centre; NIHR Biomedical Research Centre; MRC Health Data Research UK (HDR UK) Midlands; Computer, Electrical and Mathematical Sciences & Engineering Division, Computational Bioscience Research Center, King Abdullah University of Science and Technology; University Hospitals Birmingham NHS Foundation Trust, Edgbaston, Birmingham, UK

**Keywords:** text mining, negation detection, context disambiguation, clinical information extraction

## Abstract

**Background:** Negation detection is an important task in biomedical text mining. Particularly in clinical settings, it is of critical importance to determine whether findings mentioned in text are present or absent. Rule-based negation detection algorithms are a common approach to the task, and more recent investigations have resulted in the development of rule-based systems utilising the rich grammatical information afforded by typed dependency graphs. However, interacting with these complex representations inevitably necessitates complex rules, which are time-consuming to develop and do not generalise well. We hypothesise that a heuristic approach to determining negation via dependency graphs could offer a powerful alternative.

**Results:** We describe and implement an algorithm for negation detection based on grammatical distance from a negatory construct in a typed dependency graph. To evaluate the algorithm, we develop two testing corpora comprised of sentences of clinical text extracted from the MIMIC-III database and documents related to hypertrophic cardiomyopathy patients routinely collected at University Hospitals Birmingham NHS trust. Gold-standard validation datasets were built by a combination of human annotation and examination of algorithm error. Finally, we compare the performance of our approach with four other rule-based algorithms on both gold-standard corpora.

**Conclusions:** The presented algorithm exhibits the best performance by f-measure over the MIMIC-III dataset, and a similar performance to the syntactic negation detection systems over the HCM dataset. It is also the fastest of the dependency-based negation systems explored in this study. Our results show that while a single heuristic approach to dependency-based negation detection is ignorant to certain advanced cases, it nevertheless forms a powerful and stable method, requiring minimal training and adaptation between datasets. As such, it could present a drop-in replacement or augmentation for many-rule negation approaches in clinical text-mining pipelines, particularly for cases where adaptation and rule development is not required or possible.

## 1 Introduction

A major component of information extraction pipelines are algorithms that determine the context of mentioned entities. It is only with information concerning the context with which an entity has been mentioned, that the overall relationship between an object and subject can be discerned. This is a critical component of the relation extraction process. For example, a mention of a disease in a clinical letter does not necessarily imply that a patient suffers from that disease. Since many clinical letters discuss a diagnostic process, a letter may discuss a test being conducted to determine whether a patient has a condition, or refer to discussion of the arguments for whether or not a patient has a condition. The letter may also rule out the condition, or only mention that the condition is present in a family member.

In clinical text mining, one particularly important context disambiguation task is negation detection. Negation detection determines whether a clinical finding mentioned in a narrative is present or absent, usually using the sentence mentioning the concept as input [1]. Many methodologies have been applied to the task of negation detection, traditionally using rule-based methods, with a more recent focus on machine learning algorithms. The performance of rule-based methods for negation in comparison to machine learning methods is hotly contested. Two independent reviews comparing rule-based and trained machine learning approaches reported rule-based approaches to be superior in performance [2,3]. Another work reported that machine learning models modestly outperformed rule-based classifiers with out-of-context training, yielding further improvements with in-context training [4]. However, this work compares several machine learning classifiers with one particular implementation of NegEx, that is not used in any of the previous two studies, revealing superior rule-based performance, and does not consider it for additional training in any context (e.g. rule development). Another ML negation work evaluated several machine learning methods, but did not consider any rule-based methods, even though the performance was similar to rule-based methods presented elsewhere in the literature [5].

Unlike machine learning models, rule-based approaches employ classifications resulting from the application of one or more rules that are understandable from the point of view of a human operator. While more recent approaches, such as NegBERT [6] can generate an explanation for an individual case of negation, the underlying decision model cannot easily be interpreted or modified, while many other machine learning approaches cannot generate explanations at all. The ability to understand, examine, and modify the decision model surrounding context disambiguation classifiers is important in the context of any application to clinical decision making. This constitutes both a practical and an ethical concern.

NegEx is an early example of a negation detection algorithm used in the clinical domain [7], using rules described by regular expressions that make decisions based on the presence and position of tokens appearing in sentences. This approach was later generalised into ConText [8], and some form of NegEx implementation is a frequent component of modern clinical text mining pipelines, such as CogStack [9] and cTAKES [10]. These approaches can be categorised as syntactic rule-based negation detection algorithms, and measuring distance between tokens is known as a hotspot approach.

A more recent development for rule-based algorithms is the inclusion of decision models that operate upon grammatical sentence models and word relations, rather than the word tokens themselves. In particularly, these approaches utilise the typed dependency graph produced by the dependency resolution task. Dependency resolution uses a transition-based parser to construct a graph of governing and dependent word relationships in a sentence. The hypothesis of such work is that typed dependency rules should enable greater discernment, attuned to grammatical nuance beyond the mere mention of a word or appearance of a symbol in a text.

One such approach, DEEPEN [11], operates upon concepts that NegEx determines to be negated [11]. Other dependency-based algorithms make no use of NegEx, such as NegBio, negation-detection, and DepNeg [12–14], reported to exhibit an improved precision over syntactic approaches. However, an independent assessment showed that ConText maintained its performance over a novel dataset, while the other approaches did not [2]. Approaches such as SynNeg have attempted to extend ConText with more specific grammatical rules, but showed only modest performance improvements [15].

Therefore, while dependency resolution methods have proven powerful, and the extra information they afford can lead to improved performance over the corpus upon which they are trained, it seems that grammatical rules do not generalise as well as syntactic rules, while also necessitating more expert knowledge input and time to develop. The inherent complexity and ambiguity of human language leads to such a variety of grammatical models for sentences that no satisfactory set of rules can be determined via manual curation over a small set of sentences.

We hypothesise that a single-rule approach to typed dependency negation detection will perform and generalise better than more expressive rule-based approaches, leveraging particular grammatical relationships. We propose an algorithm that avoids defining specific patterns of dependency, and instead determines a measure of grammatical distance and relatedness in the dependency graph. We anticipate that such an approach will require minimal training, and provide consistent high performance across clinical datasets.

In this paper, we present a novel negation detection algorithm and compare its performance with a number of other rule-based negation detection algorithms, both syntactic and dependency-based, over two medical corpora. To this end, we develop two gold standard negation detection corpora using text sampled from the MIMIC-III critical care database [16], and routinely collected clinical letters at University Hospitals Birmingham (UHB), mentioning hypertrophic cardiomyopathy (HCM).

## 2 Materials and Methods

The negation algorithm is implemented in Groovy, making use of the transition-based neural network implementation of dependency resolution included in the Stanford CoreNLP suite [17]. CoreNLP was also used to annotate the test corpus, making use of the RegexNER annotator with all non-obsolete subclasses of *Phenotypic abnormality* (HP:0000118) in the Human Phenotype Ontology (HP) [18]. These components were used via the Komenti semantic text-mining tool, available at http://github.com/reality/komenti.

### 2.1 Algorithms

The negation detection implementation presented is named komenti-negation: the negation detection module of the Komenti text-mining framework. The algorithms we selected to evaluate against komenti-negation algorithm were NegEx, pyConTextNLP, negation-detection, and NegBio. The sources of the algorithms and version numbers used are listed in Table 1. For the reasons discussed in the introduction we only consider rule-based algorithms. The first two use syntactic rule-based negation, and the latter two utilise dependency resolution. Of the two dependency-based negation algorithms, NegBio relies on a multitude of specific grammatical rules, while negation-detection is another example of a heuristic approach.

**Table 1.** The negation algorithms considered for comparison with komenti-negation, including versions and source for download.

| Name | Type | Version | Source |
| --- | --- | --- | --- |
| NegEx | Syntactic | 21b013c | https://github.com/chapmanbe/negex/ |
| pyConTextNLP | Syntactic | 0.7.0.1 | https://pypi.org/project/pyConTextNLP/ |
| negation-detection | Dependency | 6d9d88e | https://github.com/gkotsis/negation-detection |
| NegBio | Dependency | d025875 | https://github.com/ncbi-nlp/NegBio/ |

Syntactic classifiers define a set of regular expressions that define a negatory construct. For example, PyConTextNLP includes the following rule:

Comments : ‘’
Direction : forward
Lex: without sign
Regex : without sign (s)?
Type : DEFINITE_EGATED_EXISTENCE

In the case that a **Regex** property is defined, a regular expression is used to match against the sentence, while the **Lex** property is used to define a particular phrase. The **Direction** property stipulates whether the negatory construct (pattern or absolute phrase) should appear before or after the concept. The algorithm determines internally how far the concept should be from the negatory construct in the sentence. A sentence is classified for negation depending on whether it matches one of these rules.

While syntactic rules match against the actual text of a sentence, dependency-based algorithms use rules that are matched against a graph of typed dependency relationships between words in a sentence. This usually involves determining the relationship between a node matching a vocabulary of negating words and the concept of interest. One common source of error for dependency resolution algorithms is that they cannot operate if the dependency resolution process cannot construct a model of the sentence. Most algorithms produce a default result of ‘not negated’ in this case. However, most dependency parsers can produce dependency models for grammatical sub-components of sentences.

The negation-detection algorithm is the closest to our proposed algorithm, in the sense that it repeatedly applies a single subgraph selection rule to the typed dependency graph to prune irrelevant clauses and intermediate nodes from a sentence, before employing what is essentially a string search for negation vocabulary [13]. It is reported as having a similar performance to ConText, with a slightly higher recall. In an independent assessment [19], it was reported to perform extremely well when extended with a richer negation vocabulary [19], outperforming other popular methods.

### 2.2 Corpus Generation and Training

The MIMIC dataset was derived from the MIMIC-III critical care database [16]. Entries were sampled randomly from the NOTEEVENTS table, of which one sentence was randomly selected. 500 sentences were selected for training in this way. We used Komenti to annotate the 500 sentences with biomedical concepts, running the negation algorithms against the set, and examining error cases to identify additional negatory vocabulary not currently included in the software, and to identify any errors in the test implementation, counting algorithms, or algorithms themselves. When additional negatory constructs were identified, all negation detection implementations were extended with them (if not already present). In the case of NegBio, only grammatical rules were accepted. Therefore, for each of the two negatory words that were missing: ‘deny’ and ‘not’, a bi-directional rule was introduced. It is possible that more finely tuned rules could have been developed to achieve a better performance, but the only training considered in this experiment was the addition of extra negatory words, rather than rule development. After training, a further 7,000 sentences were sampled from MIMIC for testing. These were annotated with HPO terms using the Komenti tool, yielding 1,300 annotations. HPO query and sampling were both performed on 28/12/2019.

Another validation was performed on clinical letters routinely collected at University Hospitals Birmingham (UHB), mentioning hypertrophic cardiomyopathy (HCM). 5,000 sentences were sampled from a pre-existing clinical text corpus of documents containing HCM keywords. To sample the corpus, a file was selected at random, and then one sentence was selected randomly from that file. The sentences were annotated with HPO terms using the Komenti tool, yielding 1,077 annotations. To assess algorithm generalisability and performance on an unseen dataset, no training set was used for the HCM validation, although modifications made in the MIMIC training were included in the models.

In both investigations, sentence selection criteria were used to constrain the text returned from the corpus. This served two purposes. First, to ensure that narrative text was returned, rather than field-based, table-based, or irrelevant text. Second, to limit the length of sentences so as to make it easier to perform manual validation. Sentences shorter than 4 words and longer than 30 words were excluded, and sentences containing phrases indicating field data were excluded. Sentences with indicators of unusable content (e.g. due to scanned documents) were also removed. These problems could be solved by additional pre-processing of the text, but this task is not the subject of this investigation, and using shorter sentences should not advantage any particular algorithm (although the dependency parsing algorithms are more sensitive to correct grammar). These parameters and pre-processing options were manually tuned, and decided during the training phase. For simplicity, where a single concept was mentioned multiple times in a single sentence, only one annotation was preserved, and negated concepts were given priority. This is potentially a small source of error, but should not favour any particular algorithm. The test code was designed to ask, in each case, “is an instance of the word negated within this sentence?”

In both cases, all annotations were manually labelled with respect to their negation status, determining whether the annotated concept was negated in the sentence. Concepts were marked as negated if the sentence expressed that the finding was absent. This is due to the purpose of the negation detection algorithm, in this context, being the exclusion of concept mentions from evidence of a patient having a condition if they do not have it.

### 2.3 Evaluation

We sought to make a gold standard dataset with which to make future negation algorithm comparisons. To do this, we examined the errors (false positives and false negatives) of the three best-performing algorithms by f-measure, deciding upon labels that were incorrect. We then repeated the validation. The presented results are from the corrected dataset.

The Linux command *time* was used to measure the execution time for each algorithm, with the actual elapsed time measurement taken. Apart from NegBio, two separate machines were used for the MIMIC and HCM investigations. NegBio did not run on the machine used for the MIMIC evaluation, and therefore the same machine used for HCM was used. We measured the performance of the algorithms using precision, recall, and f-measure (F1 score).

## 3 Results and Discussion

The algorithm implementation is available as part of the Komenti semantic text-mining tool, which is freely available under an open source licence at http://github.com/reality/komenti.

### 3.1 Algorithm

In our approach, rather then using a set of dependency rules, we use a measure of ‘dependency distance,’ the distance in a typed dependency graph between a negatory construct and the target term, as the measure of negation context. Therefore, we mirror the generic and transferable ‘hotspot’ methods employed by NegEx and ConText, while extending them with the notion of grammatical relatedness afforded by dependency models. In addition, because the dependency resolution process is run once, rather than at every stage of pruning, it should be faster than methods that rely on repeated reclassification of the sentence.

Algorithm 1 describes the komenti-negation algorithm, which determines whether a concept is negated in a sentence. The dependency resolution algorithm produces a typed dependency graph from a sentence, which is passed to the algorithm as input. This graph is formed of nodes that represent word tokens, and edges that represent their grammatical relationships. Together, they form a grammatical model of the sentence. Each edge is labeled with a particular kind of relationship, such as negation or adjectival noun modification. The edges also have a direction, that define a **governer** and **dependent** for each relationship. For example, in a noun modification relationship between the words **light**and **touch**, the governer would be **touch**, and the dependent **light**, because the subject is the noun **touch**, while **light** is its modifier. The graph can also be thought of as a set of assertion triples: a governer (subject), dependent (object), and relationship (predicate). Two examples of typed dependency graphs are shown in Figure 1.

**Algorithm 1:**
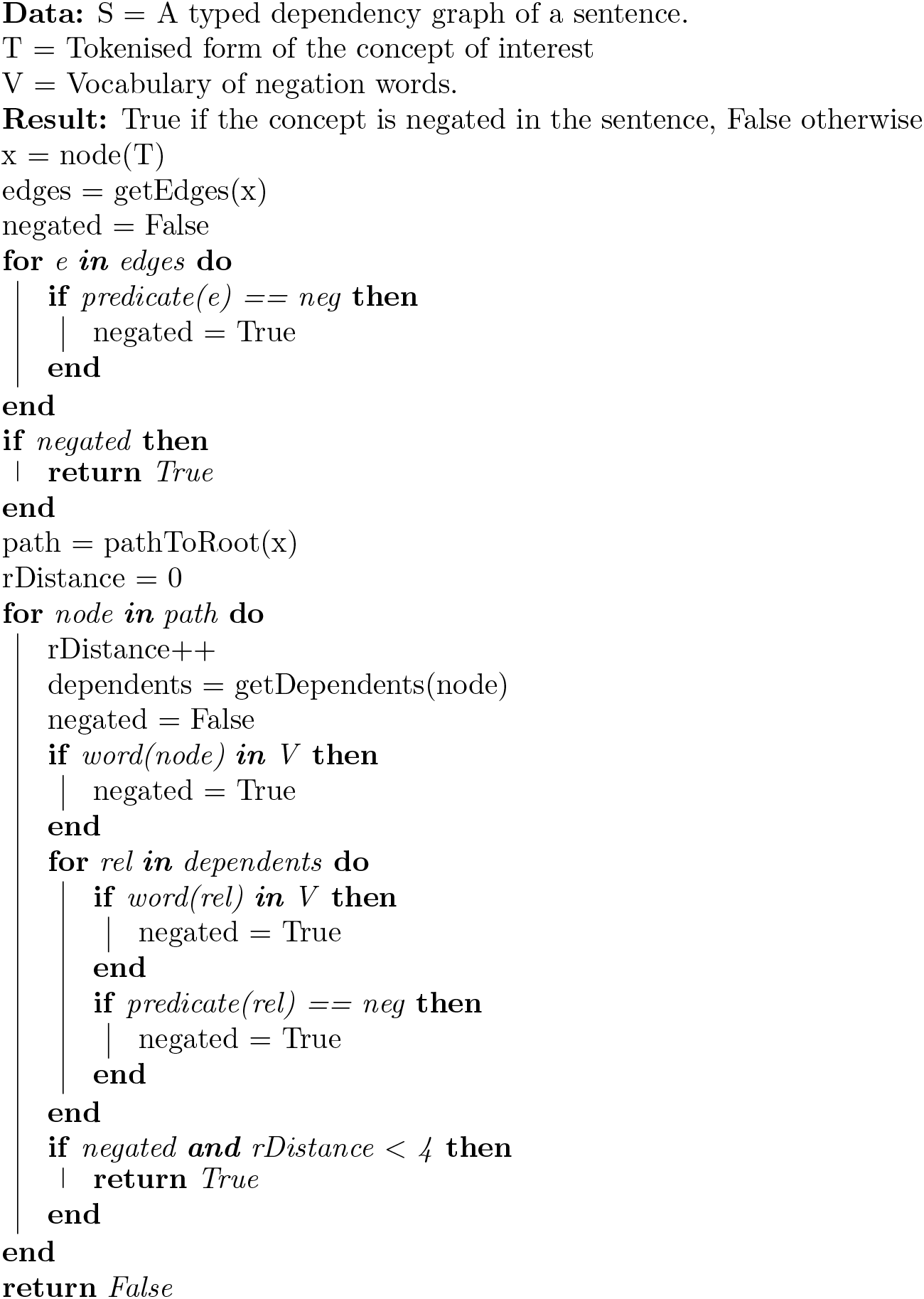
komenti-negation algorithm for determining the negation of a concept in a sentence.

**Figure 1.**
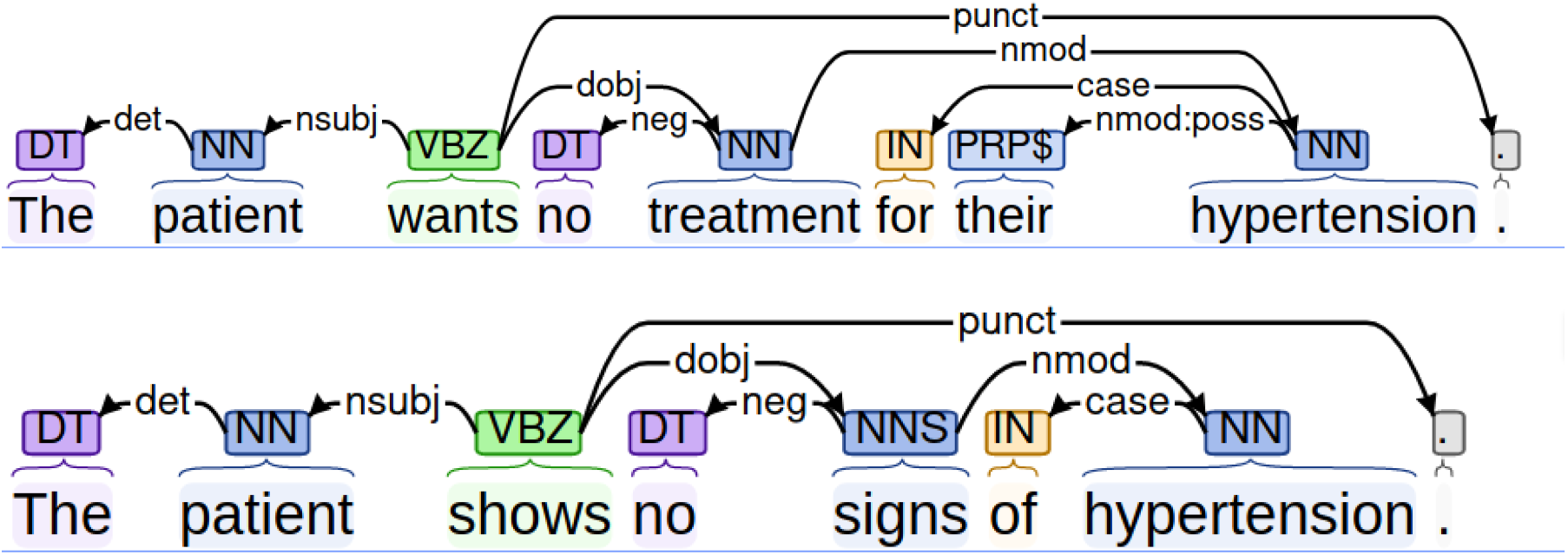
Two sentences concerning hypertension, with typed dependency annotations. The graph is formed of directional relationships between governing and dependent words, and a predicate describing the nature of the relationship between them.

Before generating the typed dependency graph, the sentence is pre-processed. Because the typed dependency graph does not support multi-word nodes, concepts of interest are replaced with a single neutral word, for which negation is then checked. The choice of this word may affect the outcome of the algorithm, due to the different dependency models they may produce. For our experiments we used ‘biscuit,’ as this is a simple and common word that does not appear in any of the considered text. In addition, if the sentence contains one of the words in the negation vocabulary, followed directly by the concept of interest, the sentence is transformed into its own sentence with the word ‘excludes’ appearing directly before the concept. This is because there is a tendency for the CoreNLP to parse such constructs into noun phrases, which does not always fall into the negated dependent model required by the algorithm. Furthermore, the transformation also makes the dependency resolution process faster.

The basis of the algorithm is an attempt to find a transitive relationship between a negation construct and the tokenised form of the concept of interest. This is done by finding the path between the concept and the root node, examining each node for a descendent relationship with either a negation predicate or object contained in the negation vocabulary. A negation vocabulary is used in addition to negatory relationships because the dependency resolution algorithm does not reliably represent all negatory constructs with a negation dependency (for example, the word ‘exclude’). Other subgraphs are not examined, as the hypothesis is that negatory constructs appearing here refer to other objects described by the sentence. If a match is found, its distance from the target concept is then measured. This relationship distance heuristic is used to eliminate unrelated negatory constructs that refer to other words, on account of their distance from the concept of interest. The maximum distance parameter can be modified, but we use a value of 4 for this experiment. This value was manually chosen during the algorithm development process.

If we take the first example from Figure 1, and evaluate the sentence “The patient shows no signs of hypertension” for negation of the ‘hypertension’ concept, we would start from the hypertension node, and move along the graph towards the root node ‘shows,’ examining the intermediate ‘treatment’ node on the way. At each step, we examine the direct descendants of the node, searching for a relationship that meets the criteria of a negatory construct, in the form of either a negation predicate, or an object that appears in our negation vocabulary. For ‘hypertension,’ we would examine the hypertension:case:of relation, which does not meet the criteria. Then we move onto the ‘signs’ node, which has two direct descendants: signs:nmod:hypertension and signs:neg:no, and the latter is a negatory construct. Since this is the second node we have examined, the distance is 2, which is below the maximum distance parameter setting of four. Therefore, the algorithm would confirm that the sentence negates the concept.

### 3.2 Evaluation

Table 2 summarises the result metrics for both the HCM and MIMIC datasets. In the MIMIC dataset, the komenti-negation algorithm performed the best by a large margin when compared by f-measure, although NegBio and NegEx had higher precision and recall respectively. In the HCM dataset, all algorithms had a better performance than the one they exhibited over the MIMIC dataset, and the disparity between algorithms was relatively small (excepting NegBio). However, NegEx performed the best on this dataset when measured by precision and f-measure.

**Table 2.** Performance comparison of negation algorithms on sentences sampled from MIMIC and HCM datasets. The best performance for each metric in each dataset is emphasised.

| Corpus | Algorithm | Precision | Recall | f-measure | Time |
| --- | --- | --- | --- | --- | --- |
| MIMIC | NegEx | 0.642 | **0.966** | 0.771 | 0m1.812s |
|  | pyConTextNLP | 0.463 | 0.936 | 0.620 | 0m53.082s |
|  | negation-detection | 0.59 | 0.66 | 0.621 | 77m56.205s |
|  | NegBio | **0.86** | 0.59 | 0.7 | 33m29.556s |
|  | komenti-negation | 0.83 | 0.91 | **0.87** | 1m10.291s |
| HCM | NegEx | **0.921** | 0.911 | **0.916** | 0m0.987s |
|  | pyConTextNLP | 0.86 | **0.932** | 0.895 | 0m41.701s |
|  | negation-detection | 0.893 | 0.874 | 0.884 | 4m26.738s |
|  | NegBio | 0.72 | 0.632 | 0.672 | 23m43.932s |
|  | komenti-negation | 0.881 | 0.89 | 0.885 | 1m16.265s |

With respect to generalisability, the Komenti algorithm maintained a stable performance over both datasets. It has, by far, the smallest magnitude of difference in f-measure between the two investigations, of 0.015, followed by NegBio with 0.028.

The performance metrics also confirmed that dependency-based negation detection systems while highly precise, they lack rules able to cover most negatory constructs when applied to novel datasets. However, this effect is not so obvious in the HCM dataset case, where the precision is quite low compared to other methods. We hypothesise that this is due to the HCM dataset containing a motif that confounds one of its developed rules. One reason that NegEx consistently performs so well across independent investigations is that its set of triggers are extremely complete and mature. It’s possible that with sufficient training approaches, such as NegBio, could achieve similar levels of maturity, including explication of rules for the handling of edge-cases.

#### 3.2.1 Running time

The quickest algorithm in both cases was NegEx, finishing in less than 2 seconds. pyConText finished in less than a minute in both cases. These are both the syntactic rule-based negation detection systems, utilising regular expressions. A single regular expression match can be found in *O*(*n*), and so the overall running time is *O*(*mn*), where *n* is the length of the sentence and *m* is the number of rules.

Meanwhile, the running time of the dependency parser algorithms is quite variable. All make use of the Stanford CoreNLP dependency resolution implementation. This employs the arc-standard system [20], which can construct a parse with a linear time treatment of each word. However, the negation-detection algorithm works by examining the graph to prune irrelevant components, and then runs the dependency resolution process again, repeating this process until it identifies what it considers to be the minimal clause surrounding the concept of interest. It should be noted that the Stanford CoreNLP program requires a number of other processes to be executed prior to dependency resolution, including tokenisation and part-of-speech tagging.

In the case of NegBio, the dependencies are only parsed once for each sentence. Each grammatical rule is converted into a graph, and then matched against the typed dependency graph using a subgraph matching algorithm. The subgraph matching algorithm is *O(m^2^k^m^)* where m is the length of the input, and k is the vertex degree, and must be repeated for every rule [12].

Komenti also uses the dependency resolution algorithm only once per sentence, and does not use an expensive sub-graph matching algorithm. The algorithm works step-by-step from the node representing the concept of interest to the root. In the worst case this is equal to the depth of the parse tree, which is *n* in its worst case (where *n* is the number of words), and *log*(*n*) on average. At each step, the direct dependents of the current node are examined (to check both its predicate status, and token content of the linked node). In the worst case, the overall number of dependent nodes examined cannot exceed *n* − 1, in which case the distance between the node representing the concept of interest and the negatory construct cannot exceed 1 (every node other than root in the sentence being a dependent of the concept of interest or the root node). Since the number of these examinations is co-dependent, it would be better to characterise the algorithm in terms of the total number of nodes examined, combining the overall iteration and descendent examination steps. At each step along the path towards the root, the descendants will include the previously examined node. Therefore, the worst case is the one where every node exists on the path to root, causing every node but the root to be examined twice, leading to a time complexity of *O*(2*n* − 1).

Both regex-based systems are extremely fast, while the Komenti algorithm finishes in just over a minute for the slower case. A discussion of memory and time usage in CoreNLP notes that there is a ‘warm-up’ period for the software, and that document processing speed can expect to increase after an initial period [21]. Since our experiments concern a relatively small number of sentences, it’s possible that a large component of Komenti’s running time is initialisation, and that processing a greater number of sentences would see an improvement in running time with respect to size of input.

However, it is shown that the repeated annotation and dependency resolution leveraged by negation-detection is a serious limitation on running time. The implementation includes log output for the CoreNLP system, and while processing the 1,077-sentence HCM dataset, it reported that more than 5,000 documents (pieces of text) were processed, at a rate of 19.839 documents per second. Meanwhile, NegBio is slow in both cases, in the faster case taking over 23 minutes to parse 1,077 sentences. We expect that this is caused by the expensive sub-graph matching algorithm.

#### 3.2.2 Sources of error and limitations

A striking feature of the results is the drastic difference in performance exhibited by most algorithms between the MIMIC and HCM datasets. While komenti-negation maintains relatively stable performance over both, average performance was much worse over the MIMIC dataset, despite the fact that it was considered for training in the form of adding additional negation vocabulary.

Through examination of the errors, we discovered that the MIMIC dataset contained much more complicated clinical descriptions of patients, which were more likely to confound the provisions made by most negation detection systems. It also contained more cases of ill-formed and ungrammatical text, in the form of run-on sentences, lack of punctuation, and text extracted without formatting from forms and tables. We attempted to limit this in our pre-processing steps, and effects from these features could be further mitigated in this way. For example, by transforming sentences that appear like fields or tables, or detecting unmarked sentence splits. Indeed, some of these processes are regular features of text mining pipelines, but are not tasks under the remit of the negation detection systems themselves. It may be considered for future work to explicitly investigate the effects of differing levels of pre-processing and sanitisation on performance of negation detection methods.

The negatory word ‘resolved’ appeared in the MIMIC test set, although it did not appear in the training set or the HCM set at all. Also, since MIMIC discusses critical care cases, it includes a greater variety of language, discussing phenotypes in a more short-term sense. In this way, it was more likely for a condition that the patient presented with but no longer had, to be discussed at length and in a complicated manner. For example, in the critical care setting described by MIMIC, people are put on a “stroke pathway” if they are deemed at risk of stroke.

Sentences with uncommon or ambiguous negation status depending on the attached noun were common: “the patient presented with chest pain, which has since been controlled” - a formulation that none of the negation detection algorithms would be able to identify without modification. Furthermore, if this were concerning if this was applied to hypertension it would be ‘treated’ hypertension, but they would still have it. To our knowledge, no negation detection system currently contains rules that examine the content of the object considered for negation status (although it would theoretically be possible for any rule-based system). This example also demonstrates another variable problem at the core of clinical negation detection, namely the differing definition of negation depending on the interests of the study. For example, a text mining project investigating critical care patients may have different definitions for the presence or absence of a concept depending on whether or not they are interested in conditions that the patient still had at discharge, or had historically but have since been resolved. In this investigation, we focused on the presence and absence of the concept local to the sentence itself, working with the clinical expert’s intuitive notion of negation. Nevertheless, this led to some ambiguity. For example, one sentence discussed a patient having had a cataract removed, and was not marked as negated, because the presence of the cataract at all was being considered as a piece of evidence for the presence of a cardiac disease. Meanwhile, several sentences that discussed patients who had been made “pain free” were marked as negated. Ultimately, some of these problems can be resolved by the integration of negation detection results into the context of larger pipelines which also discern information concerning temporal status and bearer of the findings being analysed.

We also discovered a bug in both the NegBio and negation-detection implementations, wherein they mishandle parenthetical text, causing them to be ignored, and never be marked positive for negation. Forms such as “(not shock)” or “(not delirium)” were very common in the MIMIC dataset, yet did not appear at all in the HCM texts.

We believe that part of the reason for the success of the komenti-negation approach in this context is its sensitivity to the negation dependency type, which allows for negatory constructs, not explicitly coded, to be identified, based on the overall training of the dependency resolution system. Furthermore, komenti-negation is more robust to ungrammatical sentences, because the dependency parser can isolate grammatical components of ungrammatical texts provided to it. For the same reasons, we believe that the negation-detection algorithm would reach a similar level of performance over the MIMIC dataset, if the problem of parenthetical text treatment was fixed.

Another frequent source of error, mostly endemic to the dependency resolution approaches, was the semantic effect of intermediate words on negation status. Figure 1 shows two sentences concerning a patient’s relationship with hypertension. In both cases, the pattern is very similar, in the sense that there is a transitive relationship between ‘hypertension’ and a negated noun. These nouns are ‘treatment’ and ‘signs’ respectively, and they are connected to hypertension by a noun modification relationship. The grammatical relationship between the negator and the concept of interest is the same, but they express very different things. The first refers to a treatment of hypertension, the negation of which does not indicate that the patient does not have hypertension (rather the opposite), while the second refers to signs of hypertension, which if negated also indicates there is no hypertension. This difference can also depend on the verb used, and many other potential expressible constructs.

This problem can not be easily addressed. The koment-negation and negation-detection algorithms do not currently support this kind of relationship, while NegBio only understands codes for the predicates themselves (so it could not tell the difference between a ‘sign’ and a ‘treatment’). These algorithms could be modified to accept patterns of different intermediary nouns, indicating whether or not its negation also negates the target concept, and indeed these could be implemented using NegEx regular expressions. Alternatively, a simple classifier could be trained using annotated texts to learn whether or not these negations apply transitively. However, both of these approaches would introduce a complexity and necessity for training that is besides the purpose of a simple heuristic-based algorithm.

There are some limitations inherent to Komenti’s grammatical distance approach particularly. The dependency resolution process usually tends to user the operative verb of a sentence as the root of the graph. If the sentence is particularly short, nouns that may not be negated in the sentence may be incorrectly linked to a negator. For example, in “there cough has stopped, but the hypertension continues,” both cough and hypertension would be negated. This problem could easily be solved by more advanced dependency relations. Without introducing those concepts, other methods of delimiting sentences into clauses could perhaps be utilised.

Such issues could also be approached by automatically tuning the node distance parameter. Its optimal setting potentially depends upon several features of the free text the algorithm is employed upon, including the complexity and domain of the language expressed. Moreover, heuristics within the individual sentences could be used to tune the parameter: the length of the sentence, the total depth of the target concept within the grammatical model, and the total number of noun class words in the sentence. However, the aim of this work was to develop a base method for negation using co-reference models; further development of heuristics that depend upon specific sentence structures risk suffering a large number of edge cases and an inability to easily generalise.

Another limitation of the algorithm is that it currently does not support multi-word negatory constructs. Figure 2 depicts an example of such a rule in pyConTextNLP. The Komenti algorithm cannot currently model this kind of relationship, as the grammatical tagger does not recognise ‘can rule out’ as a negatory phrase (although it can recognise some multi-word entities, or represent them as noun phrases), and the dictionary is matched against each token (word) individually. While this has not proved to be a problem for the investigation described, it would be a desirable feature for further improvement of performance and in application to datasets, where negation is expressed in more complex ways. A potential solution to this would be the use of the *entitymentions* CoreNLP plugin, which allows for the parsing of multi-word tokens. The algorithm also does not support double negation. However, this problem would be easy to solve by modifying the algorithm two detect multiple negatory constructs within a single sentence.

**Figure 2.**
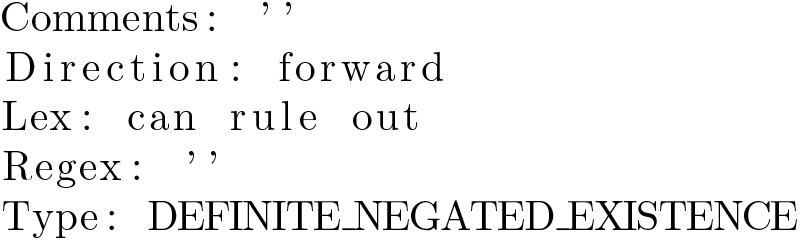
Example of a pyConTextNLP negation rule.

One potential source of dataset bias is that certain phenotypes are over represented in clinical texts, and more so in the particular kinds of clinical texts examined. This means that our evaluations do not necessarily generalise to other kinds of biomedical text. A clear example of this is ‘pain’, which accounts for 127 of the 1,300 annotations in the MIMIC dataset. Meanwhile in the HCM dataset, letters discuss people with and being investigated for HCM, meaning that HCM and its associated phenotypes are frequently discussed. In future, we could consider using a shared tasks for negation detection to evaluate the algorithm. We did not do this in this case, because our experiments were motivated and informed by the creation of a negation algorithm suitable particularly to the tasks described here, for use in a larger context with those datasets and data.

## Conclusions

Our results show that the komenti-negation algorithm was either able to out-perform or reach a similar level of performance to a number of syntactic and dependency-based negation detection algorithms, with minimal training (the addition of negation vocabulary). The algorithm maintains stable performance over two datasets, across two different dialects of English, and two different healthcare settings, namely critical care and clinical letters. In the future, we will explore its application in different settings and in particular non-biomedical ones, wherein texts often express more complicated kinds of negation.

Furthermore, the performance of the komenti-negation algorithm demonstrates that heuristic approaches applied to dependency-based negation detection enable the use of the rich information provided by typed dependencies, while remaining general enough to preserve performance across datasets and requiring minimal manual adaptation. The lack of generalisability exhibited by previous approaches to dependency-based negation is a major reason that NegEx continues to be used as a default in many clinical text mining pipelines, despite their underlying support for routines involving dependency resolution. For this reason, heuristic approaches may represent a new default choice for clinical negation detection, particularly for situations in which rule development is not possible or necessary. The komenti-negation algorithm in particular, with its high performance, low running time, and relative robustness to grammatical instability could be considered as a drop-in component for clinical text mining pipelines.

## Author’s contributions

LTS conceived of the study and experimental design, performed the experiments, implemented the software, and created the manuscript. WB and DM performed data validation. RH and GVG contributed to the manuscript. RH and GVG supervised the project. All authors revised and approved the manuscript for submission.

## Acknowledgements

GVG and LTS acknowledge support from support from the NIHR Birmingham ECMC, NIHR Birmingham SRMRC, Nanocommons H2020-EU (731032) and the NIHR Birmingham Biomedical Research Centre and the MRC HDR UK (HDRUK/CFC/01), an initiative funded by UK Research and Innovation, Department of Health and Social Care (England) and the devolved administrations, and leading medical research charities. The views expressed in this publication are those of the authors and not necessarily those of the NHS, the National Institute for Health Research, the Medical Research Council or the Department of Health.

RH and GVG were supported by funding from King Abdullah University of Science and Technology (KAUST) Office of Sponsored Research (OSR) under Award No. URF/1/3790-01-01.

## References

1. Nadeau, D., Sekine, S.: A survey of named entity recognition and classification. Lingvisticæ Investigationes 30(1), 3–26 (2007). doi:10.1075/li.30.1.03nad

2. Goryachev, S.: Implementation and Evaluation of Four Different Methods of Negation Detection, 7

3. Taggart, M., Chapman, W.W., Steinberg, B.A., Ruckel, S., Pregenzer-Wenzler, A., Du, Y., Ferraro, J., Bucher, B.T., Lloyd-Jones, D.M., Rondina, M.T., Shah, R.U.: Comparison of 2 Natural Language Processing Methods for Identification of Bleeding Among Critically Ill Patients. JAMA Network Open 1(6), 183451 (2018). doi:10.1001/jamanetworkopen.2018.3451

4. Wu, S., Miller, T., Masanz, J., Coarr, M., Halgrim, S., Carrell, D., Clark, C.: Negation’s Not Solved: Generalizability Versus Optimizability in Clinical Natural Language Processing. PLoS ONE 9(11) (2014). doi:10.1371/journal.pone.0112774

5. Taylor, S.J., Harabagiu, S.M.: The Role of a Deep-Learning Method for Negation Detection in Patient Cohort Identification from Electroencephalography Reports. AMIA Annual Symposium Proceedings 2018, 1018–1027 (2018)

6. Khandelwal, A., Sawant, S.: NegBERT: A Transfer Learning Approach for Negation Detection and Scope Resolution. arXiv:1911.04211 [cs] (2020). 1911.04211

7. Chapman, W.W., Bridewell, W., Hanbury, P., Cooper, G.F., Buchanan, B.G.: A simple algorithm for identifying negated findings and diseases in discharge summaries. Journal of Biomedical Informatics 34(5), 301–310 (2001). doi:10.1006/jbin.2001.1029

8. Harkema, H., Dowling, J.N., Thornblade, T., Chapman, W.W.: Context: An Algorithm for Determining Negation, Experiencer, and Temporal Status from Clinical Reports. Journal of biomedical informatics 42(5), 839–851 (2009). doi:10.1016/j.jbi.2009.05.002

9. Jackson, R., Kartoglu, I., Stringer, C., Gorrell, G., Roberts, A., Song, X., Wu, H., Agrawal, A., Lui, K., Groza, T., Lewsley, D., Northwood, D., Folarin, A., Stewart, R., Dobson, R.: CogStack - experiences of deploying integrated information retrieval and extraction services in a large National Health Service Foundation Trust hospital. BMC Medical Informatics and Decision Making 18(1), 47 (2018). doi:10.1186/s12911-018-0623-9

10. Savova, G.K., Masanz, J.J., Ogren, P.V., Zheng, J., Sohn, S., Kipper-Schuler, K.C., Chute, C.G.: Mayo clinical Text Analysis and Knowledge Extraction System (cTAKES): Architecture, component evaluation and applications. Journal of the American Medical Informatics Association 17(5), 507–513 (2010). doi:10.1136/jamia.2009.001560

11. Mehrabi, S., Krishnan, A., Sohn, S., Roch, A.M., Schmidt, H., Kesterson, J., Beesley, C., Dexter, P., Max Schmidt, C., Liu, H., Palakal, M.: DEEPEN: A negation detection system for clinical text incorporating dependency relation into NegEx. Journal of Biomedical Informatics 54, 213–219 (2015). doi:10.1016/j.jbi.2015.02.010

12. Peng, Y., Wang, X., Lu, L., Bagheri, M., Summers, R., Lu, Z.: NegBio: A high-performance tool for negation and uncertainty detection in radiology reports. AMIA Summits on Translational Science Proceedings 2018, 188–196 (2018)

13. Gkotsis, G., Velupillai, S., Oellrich, A., Dean, H., Liakata, M., Dutta, R.: Don’t Let Notes Be Misunderstood: A Negation Detection Method for Assessing Risk of Suicide in Mental Health Records. In: Proceedings of the Third Workshop on Computational Linguistics and Clinical Psychology, pp. 95–105. Association for Computational Linguistics, San Diego, CA, USA (2016). doi:10.18653/v1/W16-0310

14. Sohn, S., Wu, S., Chute, C.G.: Dependency Parser-based Negation Detection in Clinical Narratives. AMIA Summits on Translational Science Proceedings 2012, 1–8 (2012)

15. Tanushi, H., Dalianis, H., Duneld, M., Kvist, M., Skeppstedt, M., Velupillai, S.: Negation Scope Delimitation in Clinical Text Using Three Approaches: NegEx, PyConTextNLP and SynNeg. In: Proceedings of the 19th Nordic Conference of Computational Linguistics (NODALIDA 2013), pp. 387–397. Linköping University Electronic Press, Sweden, Oslo, Norway (2013)

16. Johnson, A.E.W., Pollard, T.J., Shen, L., Lehman, L.-w.H., Feng, M., Ghassemi, M., Moody, B., Szolovits, P., Celi, L.A., Mark, R.G.: MIMIC-III, a freely accessible critical care database. Scientific Data 3(1), 1–9 (2016). doi:10.1038/sdata.2016.35

17. Chen, D., Manning, C.: A Fast and Accurate Dependency Parser using Neural Networks. In: Proceedings of the 2014 Conference on Empirical Methods in Natural Language Processing (EMNLP), pp. 740–750. Association for Computational Linguistics, Doha, Qatar (2014). doi:10.3115/v1/D14-1082

18. Köhler, S., Doelken, S.C., Mungall, C.J., Bauer, S., Firth, H.V., Bailleul-Forestier, I., Black, G.C.M., Brown, D.L., Brudno, M., Campbell, J., FitzPatrick, D.R., Eppig, J.T., Jackson, A.P., Freson, K., Girdea, M., Helbig, I., Hurst, J.A., Jähn, J., Jackson, L.G., Kelly, A.M., Ledbetter, D.H., Mansour, S., Martin, C.L., Moss, C., Mumford, A., Ouwehand, W.H., Park, S.-M., Riggs, E.R., Scott, R.H., Sisodiya, S., Vooren, S.V., Wapner, R.J., Wilkie, A.O.M., Wright, C.F., Vulto-van Silfhout, A.T., de Leeuw, N., de Vries, B.B.A., Washingthon, N.L., Smith, C.L., Westerfield, M., Schofield, P., Ruef, B.J., Gkoutos, G.V., Haendel, M., Smedley, D., Lewis, S.E., Robinson, P.N.: The Human Phenotype Ontology project: Linking molecular biology and disease through phenotype data. Nucleic Acids Research 42(Database issue), 966–974 (2014). doi:10.1093/nar/gkt1026

19. Manimaran, J., Velmurugan, T.: Evaluation of lexicon- and syntax-based negation detection algorithms using clinical text data. Bio-Algorithms and Med-Systems 13(4), 201–213 (2017). doi:10.1515/bams-2017-0016

20. Chen, D., Manning, C.: A Fast and Accurate Dependency Parser using Neural Networks. In: Proceedings of the 2014 Conference on Empirical Methods in Natural Language Processing (EMNLP), pp. 740–750. Association for Computational Linguistics, Doha, Qatar (2014). doi:10.3115/v1/D14-1082

21. Understanding Memory and Time Usage — Stanford CoreNLP. https://stanfordnlp.github.io/CoreNLP/memory-time.html

